# The Effects of Mean Temperature and Diurnal Temperature Range on West Nile Virus Transmission

**DOI:** 10.64898/2026.08.03.742566

**Authors:** Laura C. Multini, Daniel Hartman, Britny Johnson, Adrienne Healy, Jared Skrotzki, Gregory D. Ebel, Courtney C. Murdock

**Affiliations:** Department of Entomology, Cornell University, Ithaca, New York, United States of America; Department of Microbiology, Immunology and Pathology, Colorado State University, Fort Collins, Colorado, United States of America

## Abstract

Temperature is a major driver of arbovirus transmission, and a changing climate is expected to affect this process significantly. Most experimental studies rely on constant temperatures and overlook realistic daily temperature fluctuations that impact vector-pathogen interactions. The nonlinear relationship between temperature and vector traits responsible for pathogen transmission suggests that daily fluctuating temperatures produce phenotypes important for transmission distinct from those at constant temperatures. Here, we tested whether realistic variations in mean temperature and diurnal temperature range (DTR) influence West Nile virus (WNV) transmission. We experimentally infected colonized *Culex tarsalis* with WNV-infectious blood meals and maintained them under constant and fluctuating temperature regimes. We quantified how temperature, DTR, and incubation time shaped several outcomes (infection, dissemination, infectiousness, WNV loads in mosquito saliva and mosquito survival). We also integrated survival and infectiousness probabilities to infer implications for transmission. We observed strong effects of temperature on the WNV infection dynamics, viral loads and mosquito survival. Although the probability of initial infection was similar among all temperature regimes, dissemination and infectiousness were constrained by temperature. Dissemination and infectiousness were highest at 26°C and reduced at cooler (22°C) and warmer (30°C) temperatures. Fluctuating temperatures significantly decreased infection and dissemination probabilities, while the probability of becoming infectious was driven primarily by mean temperature rather than DTR. Mosquito survival probability was temperature dependent, and fluctuating regimes increased mortality risk by 1.95-fold. Integrating mosquito survival and infectiousness dynamics demonstrated that, although at higher temperatures the extrinsic incubation period is shorter, reduced mosquito lifespan can limit viral transmission. Together, these results indicate that overall WNV transmission is affected by temperature-dependent viral dynamics and host physiological constraints. Fluctuating temperature regimes significantly delay viral dissemination within the host, especially at the optimal temperature (26°C). Measurement of infection outcomes under constant temperatures treatments may lead to over/under estimation of transmission parameters in epidemiological models, therefore integrating fluctuating temperature treatments into mechanistic models of transmission can improve estimates of when and where WNV transmission is most likely to occur.

## Introduction

Mosquito-borne viruses cause several diseases that impact human health worldwide. Dengue, Zika, Yellow Fever, and West Nile viruses threaten global public health, with billions of people currently living in areas at risk of transmission [1]. West Nile virus (WNV) is the most prevalent mosquito-borne arbovirus in the United States [2]. Since its detection in New York in 1999, WNV has rapidly expanded across North America and is now considered endemic throughout much of the United States [2–4], where it causes thousands of reported infections annually, including cases of severe neuroinvasive disease [5]. Understanding the environmental determinants of WNV transmission is essential for predicting arbovirus risk under changing climates.

Temperature is a key driver of arbovirus transmission, shaping mosquito life history, virus replication, and evolutionary dynamics [6–12]. Mechanistic modeling studies predict that WNV transmission exhibits a unimodal response to temperature, with peak transmission occurring at intermediate temperatures between approximately 23°C and 26°C [13]. Experimental studies further demonstrate that temperature significantly influences WNV infection, dissemination, and transmission in mosquitoes [14,15]. While several studies have demonstrated the effects of temperature on traits relevant for mosquito fitness and pathogen transmission [11,12,16,17], most experimental work has characterized these effects across a range of constant temperature conditions, neglecting the realistic daily temperature fluctuations experienced by mosquitoes in natural environments [18,19].

Fluctuating temperatures can alter arbovirus transmission in ways that are not captured by constant-temperature experiments because mosquito and virus traits respond to temperature in a nonlinear fashion [10,18]. Consistent with metabolic theory, trait performance increases from a lower limit (*T_min_*) to an optimum (*T_opt_*) due to increases in the efficiency and rates of metabolic processes, then declines approaching an upper limit (*T_max_*) due to the accumulation of physiological stress and damage as mosquitoes approach lethal temperatures [10,19,20]. Under fluctuating regimes, nonlinear averaging predicts that the effect of variation depends on whether fluctuations occur on the accelerating, peak, or declining portions of the thermal performance curve [21,22]. Fluctuations around lower temperatures may enhance performance, whereas fluctuations near upper limits may reduce performance through repeated stress exposure or, in some cases, improve performance by allowing partial recovery during cooler periods [23].

To advance our understanding of how daily temperature fluctuation influences WNV transmission, we conducted a series of laboratory experiments examining the effects of mean temperature and daily temperature fluctuation on WNV transmission by *Culex tarsalis*. Adult mosquitoes were maintained under three constant and three fluctuating temperature regimes to assess how temperature variation influences mosquito survival and WNV infection dynamics. Our results show that mosquito infection, dissemination, infectiousness, viral loads in the saliva, and mortality were highly influenced by mean temperature. While fluctuating temperatures significantly reduced infection and dissemination, their overall effect was more moderate than those reported for other mosquito-virus systems [18,24]. Collectively, our findings demonstrate the importance of incorporating realistic thermal environments into studies of mosquito-borne disease to better understand how temperature shapes arbovirus transmission dynamics, thus improving predictions of transmission risk under a changing climate.

## Results

### WNV infection dynamics in *Cx. tarsalis*

Across all treatments, we observed strong temperature-dependent effects on the probability of WNV infection, dissemination, and transmission (Fig. 1). Our GLMM analysis indicates that the probability of infection, dissemination, and becoming infectious was best explained by temperature, dpi, DTR, and the interactions between temperature and dpi. The most parsimonious model for all response variables was Model 2 (Table 1), which included linear and quadratic terms for dpi, interactions between temperature and dpi, and the main effect of DTR.

**Fig 1.**
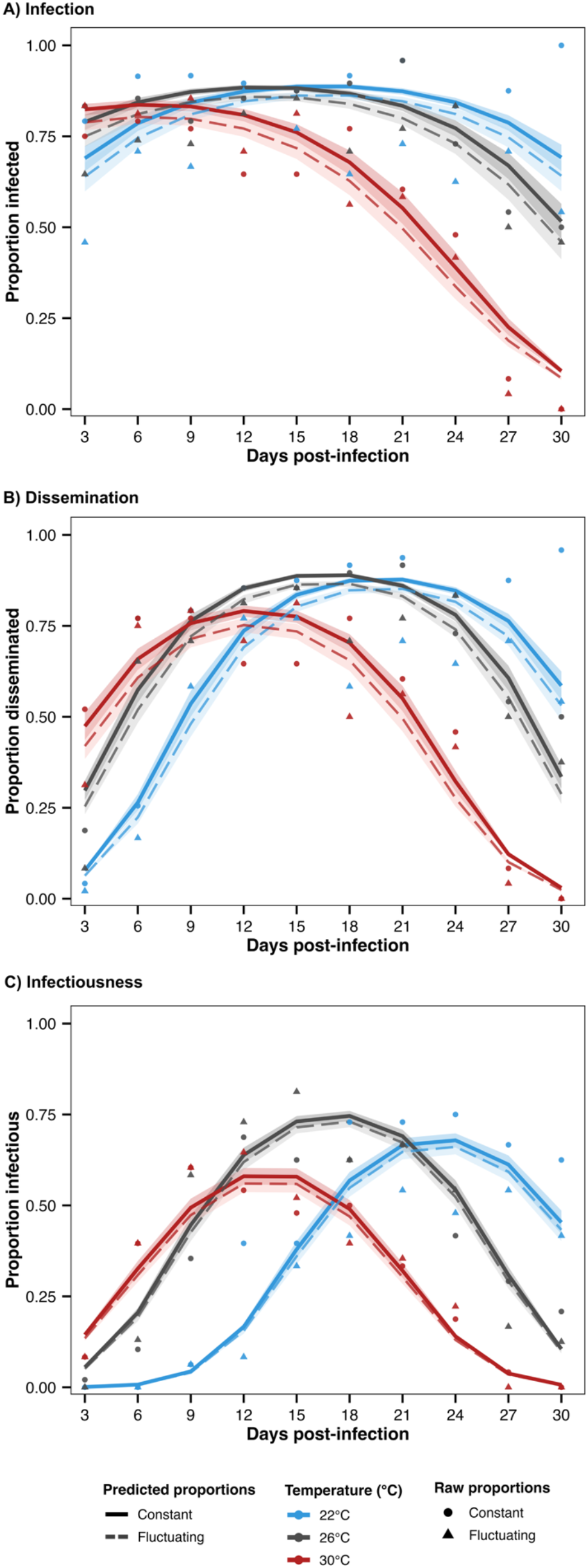
Temperature and daily temperature range shape WNV infection dynamics in *Culex tarsalis*. The relationship between days post-infection (3, 6, 9, 12, 15, 18, 21, 24, 27, 30) and the proportion of mosquitoes (A) infected (WNV-positive bodies), (B) with disseminated infections (WNV-positive legs and wings), and (C) infectious (WNV-positive saliva) out of the total mosquitoes exposed to WNV. Mosquitoes were held at three different temperatures (22°C, 26°C, 30°C) under either constant or diurnally fluctuating temperature regimes. Points represent observed proportions, shaded regions represent 95% confidence intervals around model predictions, and lines represent predicted proportions from generalized linear mixed effects models.

**Table 1.** Effects of temperature (*T*), days post-infection (*dpi*), and diurnal temperature range (*dtr*) on the probability of infection, dissemination, and infectiousness in *Culex tarsalis*. The most parsimonious generalized linear mixed model for each response variable included the terms *T x dpi*, *dtr*, and *dpi².* Models were fit using a binomial error distribution with a logit link. Coefficients are presented as log-odds estimates ± standard error (*p-value*). Mosquito cohort was included as a random effect.

| Predictor | Infection | Dissemination | Infectiousness |
| --- | --- | --- | --- |
| *Intercept* | 2.06 ± 0.37 (<0.001) | 1.63 ± 0.29 (<0.001) | -0.48 ± 0.21 (0.023) |
| *T (26°C)* | -0.04 ± 0.15 (0.783) | 0.43 ± 0.15 (0.004) | 1.48 ± 0.17 (<0.001) |
| *T(30°C)* | -0.91 ± 0.12 (<0.001) | -0.40 ± 0.12 (<0.001) | 0.79 ± 0.14 (<0.001) |
| *Dpi (scaled)* | 0.17 ± 0.08 (0.041) | 1.21 ± 0.09 (<0.001) | 2.51 ± 0.15 (<0.001) |
| *Dpi*<sup>2</sup> | -0.45 ± 0.05 (<0.001) | -1.02 ± 0.06 (<0.001) | -1.26 ± 0.07 (<0.001) |
| *DTR (fluctuating)* | -0.23 ± 0.11 (0.043) | -0.22 ± 0.11 (0.042) | -0.08 ± 0.11 (0.438) |
| *26°C × Dpi* | -0.38 ± 0.12 (0.001) | -0.79 ± 0.12 (<0.001) | -1.84 ± 0.17 (<0.001) |
| *30°C × Dpi* | -1.10 ± 0.12 (<0.001) | -1.84 ± 0.13 (<0.001) | -3.01 ± 0.20 (<0.001) |

### The effects of temperature and daily temperature range on WNV infection, dissemination, and infectiousness

Among the infection metrics measured, the probability of successful infection (WNV+ bodies) was the least sensitive to variation in mean constant temperatures (Fig. 1A). Under constant temperature regimes, infection probabilities were similar across temperatures over the course of infection. At dpi 3, the earliest time point, infection rates were already high and comparable among temperature treatments (79% at 22°C, 83% at 26°C, and 75% at 30°C). Model predictions indicated a decline in infection probabilities at higher temperatures and revealed a small but significant reduction in infection probability under fluctuating temperature regimes, particularly at cooler temperatures. Predicted infection probabilities under fluctuating conditions were 46% at 22°C, 65% at 26°C, and 83% at 30°C. Although the number of infected females declined over time at warmer temperatures, this pattern was attributable to increased mosquito mortality over the course of the experiment rather than reduced susceptibility to infection.

The probability of dissemination (WNV+ legs and wings) was highly sensitive to temperature (Fig. 1B). At 30°C, dissemination peaked early, around 12 days post-infection, and declined sharply after day 15 due to cumulative mosquito mortality. At 26°C, dissemination probabilities also peaked around day 12 but remained high until approximately day 24, before declining. In contrast, at 22°C, dissemination peaked later in the infection course, between days 18 and 21. Across all temperatures, fluctuating thermal regimes significantly reduced dissemination probabilities relative to constant temperature conditions, indicating that temperature variability constrains the progression of infection beyond the midgut.

The probability of mosquitoes becoming infectious (WNV+ saliva) was strongly temperature dependent (Fig 1C). Infectiousness was highest at 26°C, corresponding to the T*opt* for WNV transmission [13]. Model predictions showed that infectiousness peaked earlier and declined more rapidly at 30°C, whereas peak infectiousness occurred later (around day 24 post-infection) at 22°C. Unlike dissemination, mosquito infectiousness did not differ significantly between constant and fluctuating temperature treatments, suggesting that temperature variability primarily constrains earlier stages of viral progression.

### The effects of temperature and daily temperature range on mosquito survival

Temperature had a strong effect on mosquito mortality, with significantly reduced survival at both 26°C and 30°C compared to 22°C (Fig. 2; Table 2; *p* < 0.0001 for both comparisons). Mosquitoes held at 30°C experienced a more than 20-fold increase in mortality risk relative to those at 22°C (Table 2; HR = 22.03).

**Fig 2.**
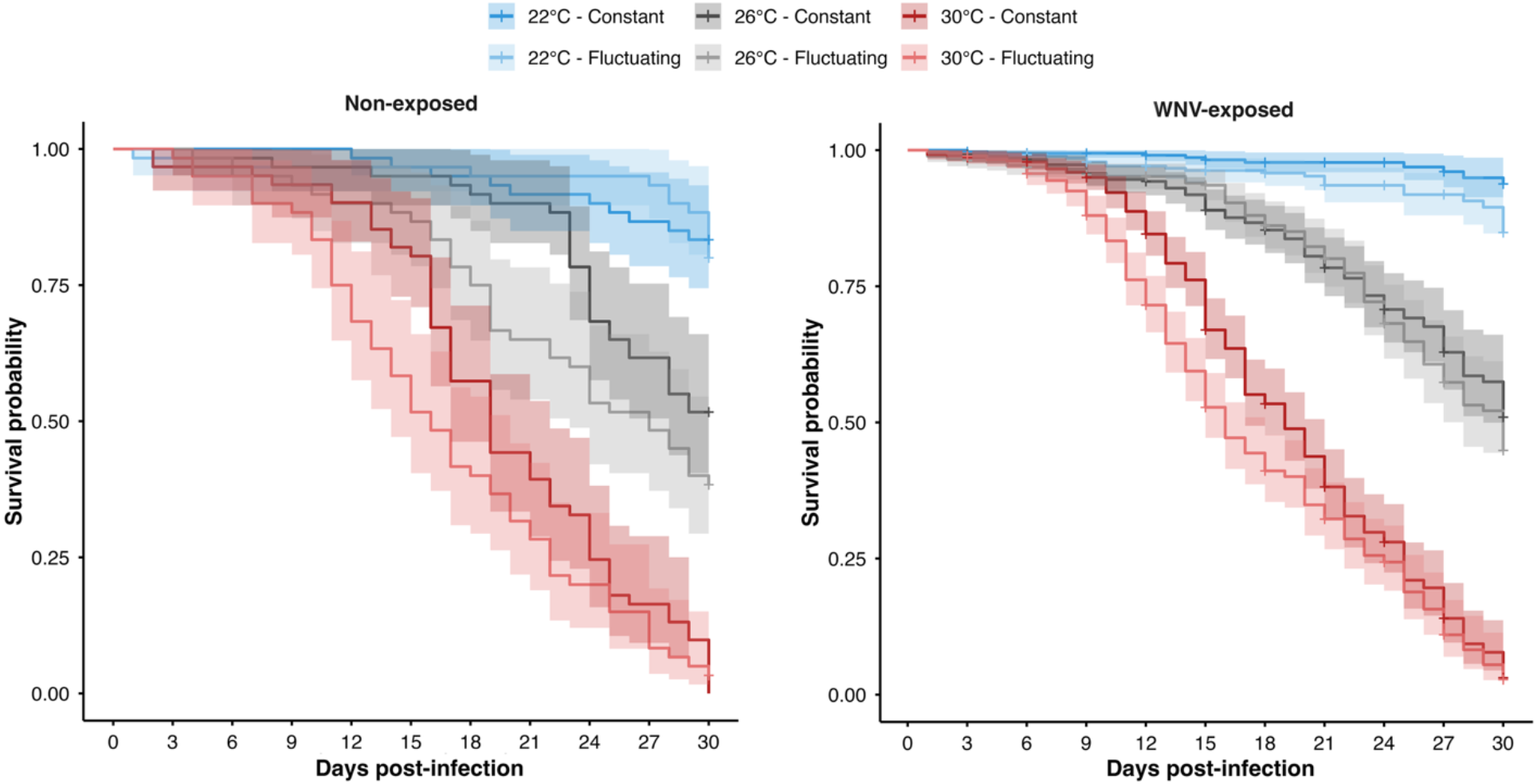
Temperature and daily temperature range influence *Culex tarsalis* survival. Kaplan-Meier survival probability curves showing mosquito survival over time under three temperatures (22°C, 26°C, 30°C) and two diurnal temperature ranges (constant and fluctuating). Survival is shown for (A) non-exposed mosquitoes and (B) WNV-exposed mosquitoes. Shaded areas indicate 95% confidence intervals.

**Table 2.** Effects of temperature, diurnal temperature range (DTR), and infection status on mosquito survival from a Cox mixed-effects model with mosquito cohort as a random effect. Reference temperature = 22°C.

| Predictor | Coefficient ($\beta$) | Hazard Ratio (HR) | SE | z | p-value |
| --- | --- | --- | --- | --- | --- |
| 26°C | 1.7549 | 5.78 | 0.3228 | 5.44 | <0.0001 |
| 30°C | 3.0926 | 22.03 | 0.3015 | 10.26 | <0.0001 |
| Fluctuating T | 0.6674 | 1.95 | 0.3043 | 2.19 | 0.028 |

Daily temperature range also significantly affected survival, with mosquitoes exposed to fluctuating temperature regimes experiencing a 1.95-fold increase in mortality risk compared to those held under constant temperatures (Table 2; *p* = 0.028). WNV infection status did not significantly influence mortality, indicating no detectable survival cost of WNV infection under the conditions tested. Interaction terms were not significant. Together, these results demonstrate that both higher temperatures and temperature fluctuations significantly reduce mosquito survival, while WNV infection itself does not contribute to mortality risk in this system.

### The effect of temperature and daily temperature range on vector competence, extrinsic incubation rate, and estimated relative force of infection

Vector competence (i.e., the maximum proportion of exposed mosquitoes that became infectious) was highest under the optimal constant temperature of 26°C (S1 Fig). Fluctuating temperature treatments decreased vector competence for both 22°C and 26°C when compared to mosquitoes housed at a similar mean constant temperature. At the highest temperature, 30°C, vector competence was low and similar for mosquitoes housed in constant and fluctuating conditions. Extrinsic incubation rate increased with temperature, showing the slowest and fastest rates at 22°C and 30°C, respectively, indicating an overall higher transmission efficiency and faster progression to infectiousness at higher temperatures (S2 Fig). Under fluctuating temperatures, EIR was higher at 26°C when compared to the constant regime, which suggests that daily temperature variation can speed infectiousness near the T*opt* for transmission. After integrating infectiousness and survival, the relative force of infection differed among temperature treatments (Fig. 3). The relative force of infection was estimated as infectious mosquito-days and expressed as the average number of mosquitoes alive and infectious per 100 WNV-exposed mosquitoes per day. This metric was calculated as the area under the curve over 30 dpi and was highest at 26°C under constant temperature. Under this condition, the cumulative infectious mosquito-days corresponded to an average of 48 mosquitoes alive and infectious per 100 WNV-exposed mosquitoes per day over 30 days. This was followed by 26°C under fluctuating regimes (avg. 46.4 mosquitoes). Relative force of infection was lower at 22°C constant (avg. 41.4 mosquitoes), and was substantially reduced at 22°C under fluctuating regimes (avg. 26 mosquitoes). At 30°C, relative force of infection was low under both constant and fluctuating regimes (avg. 29.1 and 25.7 mosquitoes, respectively).

**Fig 3.**
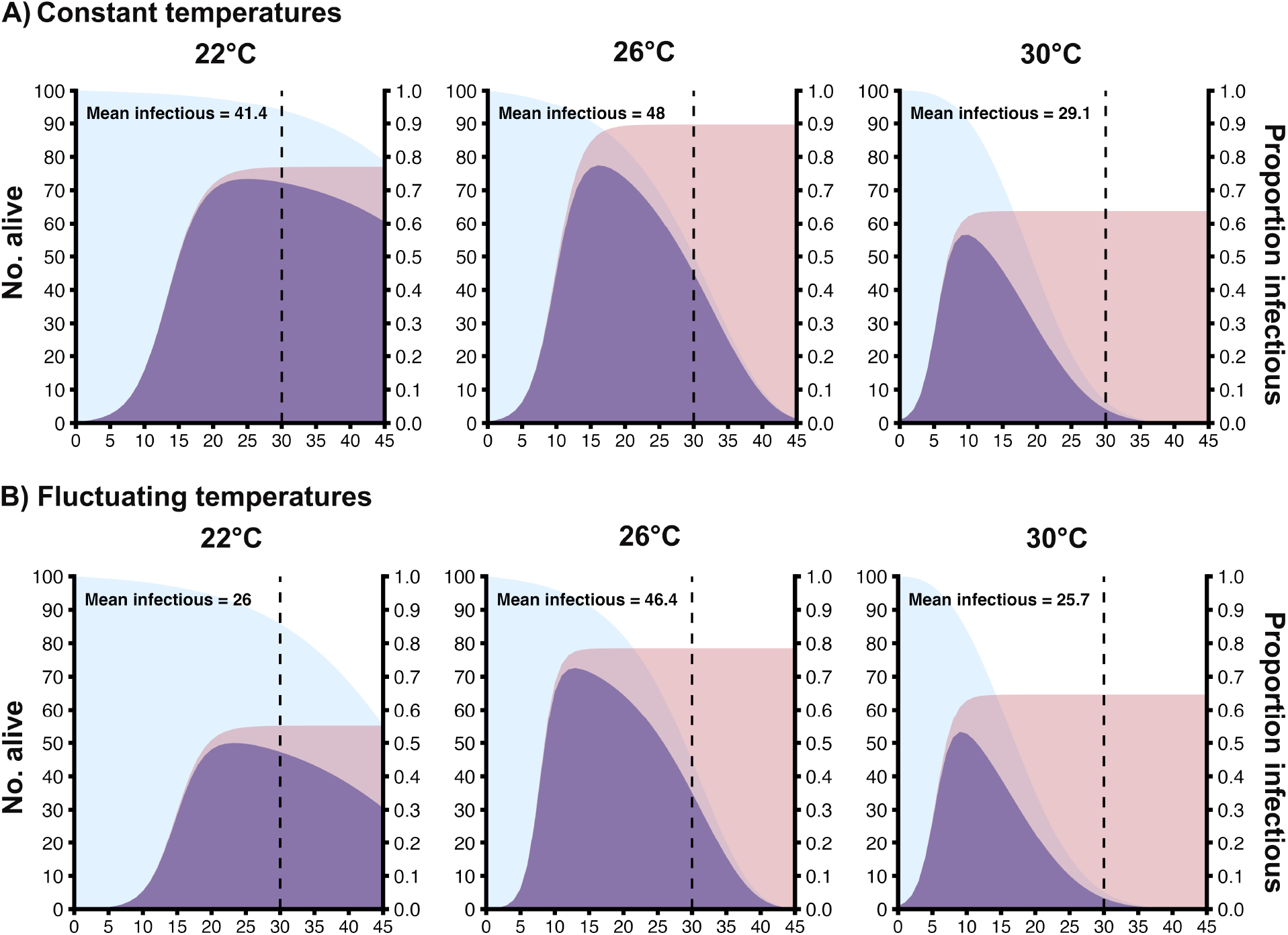
Daily proportions of mosquitoes alive, infectious, and both alive and infectious for mosquitoes exposed to WNV at different temperature regimes. Relationship between the daily proportion of mosquitoes alive (blue), infectious (pink), and those that are both alive and infectious (purple) for *Cx. tarsalis* exposed to constant (A) and fluctuating (B) temperature regimes (22°C, 26°C, 30°C). Mean infectious indicates the average number of mosquitoes alive and infectious per 100 WNV-exposed mosquitoes per day over the observed experimental period (30 days). This metric was calculated as the area under the curve. The dashed vertical line indicates the experimental endpoint at 30 dpi. Trajectories beyond day 30 are extrapolated by the model and shown to visualize predicted dynamics.

### Effects of temperature on mosquito saliva WNV viral loads

West Nile virus loads in mosquito saliva were highly variable, but the final most parsimonious linear model included significant effects of *T* and the interaction between *DTR* and *dpi* (Table 3). Viral loads were higher at 30°C than at 22°C (S3 Fig A, *p* = 0.032) and showed a significant increase with *dpi* (*p* < 0.001). Under fluctuating temperatures, the temporal increase was reduced, as indicated by a significant interaction between negative *DTR* and *dpi* (S3 Fig B; *p* = 0.013). In contrast, viral loads at 26°C did not differ from 22°C, and the main effect of *DTR* did not cause an overall shift in saliva load at the average *dpi*, but it changed how viral load varied over time (S3 Fig B).

**Table 3.**
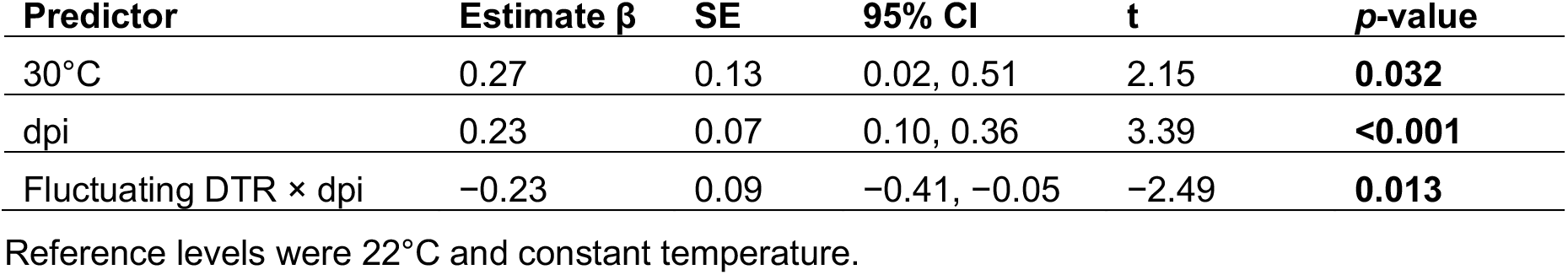
Effects of temperature, dtr, dpi, and their interaction on infectious mosquito saliva loads from a Gaussian linear model. Most parsimonious model selected included the terms *T, dtr x dpi*, and *dpi²*.

| Predictor | Estimate $\beta$ | SE | 95% CI | t | p-value |
| --- | --- | --- | --- | --- | --- |
| 30°C | 0.27 | 0.13 | 0.02, 0.51 | 2.15 | **0.032** |
| *dpi* | 0.23 | 0.07 | 0.10, 0.36 | 3.39 | **<0.001** |
| Fluctuating *DTR* x *dpi* | -0.23 | 0.09 | -0.41, -0.05 | -2.49 | **0.013** |

Reference levels were 22°C and constant temperature.

## Discussion

Land use change, increased global movement, and temperature have been identified as key drivers influencing the emergence, spread, and persistence of arboviruses [25,26]. As several studies have previously demonstrated, temperature is a key environmental driver shaping mosquito-arbovirus interactions, and its effects on arbovirus transmission are mediated by complex interactions between hosts, pathogens, and the environment [8,11,12,14,18,27–29]. However, few studies have examined how fluctuating temperatures affect pathogen transmission [18,20,30,31]. In this study, we experimentally estimated the effects of mean temperature and daily temperature range on measures of WNV transmission and mosquito mortality. Our findings demonstrate that both temperature and daily temperature range influence WNV transmission potential in *Cx. tarsalis* by shaping both infection dynamics, mosquito survival, and overall transmission potential.

Our results show that while initial infection rates were robust across temperature treatments, the daily probability of dissemination, becoming infectious, and mosquito survival were highly sensitive to temperature variation. Across all temperature treatments, midgut infection was weakly affected by temperature, with consistently high infection probabilities observed across constant temperature environments. These findings suggest that initial midgut infection represents a permissive stage in the WNV infection dynamics in *Cx. tarsalis*, consistent with previous work indicating that midgut infection is less sensitive to temperature, while downstream anatomical ‘barriers’, such as midgut escape and salivary gland infection are temperature dependent [14–16]. Dissemination was highly temperature dependent and shown to be an important thermal bottleneck in WNV mosquito infection. Midgut escape occurred rapidly at 30°C, peaking early in the infection course and declining rapidly after, a pattern driven by mosquito mortality. This dynamic has been demonstrated before and suggests that higher temperatures allow for the midgut epithelial cells to become more permeable, permitting more rapid viral escape [8,11,12,14–16]. Mechanistic models suggest that the *T_opt_* for WNV transmission by *Cx. tarsalis* is 26°C [13], which corroborates our finding that dissemination probabilities at this temperature remained high for an extended period, reflecting a balance between efficient viral replication and mosquito survival.

At 22°C, dissemination was delayed and peaked later in the infection course, suggesting slower viral replication and delayed midgut escape [32]. Although mosquitoes maintained at 22°C had high survival probabilities during the course of infection – allowing for late dissemination – such extended lifespans are unlikely under natural conditions [33]. Similar nonlinear effects of cool and warm temperatures on vector competence were observed before with *Aedes albopictus* infected with Dengue virus 2 [34], *Ae. aegypti* infected with Zika virus [12], and *Cx. pipiens* infected with WNV [14]. In contrast, previous studies demonstrated that rearing mosquitoes at cooler temperatures during the larval stage resulted in increased susceptibility to Chikungunya virus in *Ae. albopictus* [35,36]. This suggests that, although in this study we exposed mosquitoes to constant and fluctuating temperatures in the adult stage, temperature variation in both immature and adult mosquito stages are likely important in shaping arbovirus transmission [12]. The probability of becoming infectious was strongly temperature dependent and exhibited a unimodal relationship with temperature, peaking at 26°C. At this optimal temperature, mosquitoes achieved high infectiousness and survived long enough to reach the infectious stage, supporting that WNV transmission by *Cx. tarsalis* is optimized at 26°C [13].

Fluctuating temperature regimes showed a low but significant reduction in infection probability, indicating that daily thermal variability can influence early infection dynamics even when mean temperatures are permissive. A similar pattern was reported by Lambrechts et al. [18], who showed that diurnal temperature fluctuation negatively affected *Ae. aegypti* vector competence for dengue virus, which they attributed to the negative effects of both cool and high temperatures on viral progression within the mosquito [37]. Daily temperature range further constrained viral dissemination across all temperatures, with fluctuating regimes significantly reducing dissemination probabilities in relation to constant temperatures. This effect suggests that diurnal temperature fluctuation affects viral escape from the midgut and subsequent viral systemic spread, corroborating the importance of incorporating thermal variability into mechanistic models assessing arbovirus transmission risk [10,18,20].

In contrast to our other infection response variables, the probability of becoming infectious was not significantly affected by temperature fluctuations. This result suggests that temperature fluctuation primarily affects earlier stages of viral replication in *Cx. tarsalis*, such as midgut escape and dissemination to secondary tissues. However, after the salivary gland has been infected and systemic infection has been established, saliva viral loads were highly variable among individuals and dependent on temperature; mosquitoes that were held at 30°C had higher saliva viral loads than those held at 22°C. Viral loads also increased with days post infection under constant temperatures, while this increase was significantly reduced under fluctuating temperature regimes. These findings suggest that mean temperatures can influence the amount of virus expectorated in the saliva of infectious mosquitoes, while temperature fluctuations alter the viral load trajectories over time.

Based on Jensen’s inequality and the nonlinear mosquito traits response to temperature [19,23,38], we anticipated that fluctuating temperatures would reduce infection rates at the optimal temperature of 26°C and consequently limit downstream dissemination and infectiousness. At cooler temperatures, we expected that fluctuating temperatures would improve trait performance relative to constant conditions by periodically exposing mosquitoes to optimal temperatures. In contrast, at higher temperatures regimes, we hypothesized that temperature fluctuations could either improve performance of the traits tested by accelerating viral progression within the mosquito or reduce it by reaching temperatures in which mosquito mortality would constrain virus transmission. Surprisingly, fluctuating temperatures produced an overall reduction in infection and dissemination probabilities, suggesting that fluctuating temperature regimes may impose shifts in viral dynamics producing different phenotypes than expected. One possible explanation is that because WNV genome cyclization and replication initiation rates are less efficient at cooler temperatures [39], the cooler portions of the daily cycle may slow early viral replication kinetics [9,40], creating a bottleneck that is likely not fully recovered during warmer periods of the day. Another hypothesis is that fluctuating temperatures may alter host physiology, immunity, and transcriptional responses that regulate infection progression [41–44], by reducing the probability of infection and dissemination even when daily mean temperature is permissive.

Mosquito survival was significantly constrained by temperature, with warmer temperatures (26°C and 30°C) significantly increasing mosquito mortality risk relative to 22°C, and mosquitoes at 30°C experiencing more than a 20-fold increase in mortality. This pattern is consistent with the metabolic theory of ecology, which predicts that biochemical reaction rates increase exponentially with temperature up to an T*opt*, after which performance declines sharply due to protein degradation and other physiological constraints [10,45,46]. Daily temperature fluctuations significantly increased mortality risk, indicating that diurnal thermal variability imposes physiological stress on mosquitoes even when mean temperatures fall within a permissive range, likely by increasing the energetic costs of thermal acclimation and maintenance [23]. WNV infection did not significantly affect mosquito survival, suggesting that *Cx. tarsalis* experiences no survival cost of infection under the conditions tested, and the observed mortality patterns are driven by environmental temperature.

By integrating the daily probability of survival and infectiousness to estimate the relative force of infection, we demonstrate that despite the positive effects of warming temperature on the overall efficiency of infection, infection is constrained at warmer temperatures due to mosquito mortality. However, the implications for transmission are more complex at warmer temperatures. Although mosquitoes held at 30°C experienced reduced survival, they became infectious more rapidly likely due to rapid viral replication at warmer temperatures [8,9], indicating a decreased extrinsic incubation period. Because extrinsic incubation period and mosquito survival are key determinants of transmission potential, the fact that mosquitoes become infectious more rapidly can compensate for increased mosquito mortality if they are blood feeding soon after becoming infectious [10]. At cool temperatures, mosquitoes became infectious substantially later in the course of infection, likely reflecting reduced viral replication and longer extrinsic incubation period [40], which can limit transmission despite longer mosquito lifespan. Consequently, the highest relative force of infection occurred under *Topt* conditions, where mosquitoes became infectious early enough while still maintaining sufficient survival to contribute infectious mosquito-days. These results demonstrate that WNV transmission potential reflects a balance between temperature-dependent viral dynamics and vector physiological constraints.

One limitation of the study is that we tested a limited range of mean temperatures and a single magnitude of DTR (12°C). Additional cooler and warmer temperature regimes, and multiple DTR amplitudes are needed to estimate whether the effects of fluctuation become stronger near the thermal limits of WNV transmission [8,18]. Additionally, our relative force of infection estimate integrates mosquito survival and infectiousness, but does not include other temperature-sensitive trait such as biting rate [47,48]. Therefore, this metric should be interpreted as the relative availability of alive and infectious mosquitoes at a given population size rather than an absolute estimate of transmission risk. We also did not measure potential carry-over effects of different temperatures exposure, in which temperature experienced during immature stages could influence later infection, dissemination, infectiousness, and survival [27]. Finally, in order to obtain robust results and adequate sample sizes, and to control the age of our mosquitoes, we used a well established mosquito colony. It may be that the physiological responses of this colony differ from *Cx. tarsalis* in the field, which are subject to a more variable thermal environment. Nonetheless, our results provide a critical baseline for understanding how temperature influences WNV transmission by this important vector. Future studies incorporating these metrics and utilizing additional colonies or field mosquitoes, would help identify the mechanisms underlying the trends observed in this study.

Our findings highlight that incorporating both constant and fluctuating temperatures into forecasting frameworks can improve early warning systems and help target surveillance and vector control to periods when interventions are most likely to reduce the emergence and persistence of infectious vectors. Although mechanistic models based on constant temperatures may capture a substantial portion of transmission potential, incorporating temperature fluctuations may still be important when considering traits that impact transmission such as mosquito lifespan, bite rate and fecundity [19,49]. The same thermal dynamics that shape transmission potential are also likely to influence evolution and ecological dynamics of WNV transmission. Because RNA viruses are able to evolve rapidly as a response to their replicating environment through an “error-prone replication” [50], warmer temperatures may increase mutation rates and facilitate the emergence of highly adaptive WNV strains [9]. This process, in turn, drives local adaptation of WNV populations to specific thermal environments [6]. A key next step is to evaluate how temperature and daily temperature range shape the evolutionary dynamics of WNV populations during mosquito infection, dissemination and infectiousness. Together, these efforts will improve our ability to predict not only when WNV transmission is likely to occur, but also when thermal conditions may favor the emergence of more infectious or better-adapted viral populations under current and future climate conditions.

## Material and Methods

### Mosquito Husbandry

*Culex tarsalis* from a longstanding colony (KNWR – Kern National Wildlife Refuge, established by WK Reisen in 2001[51]) were obtained and maintained at Cornell University as follows. Mosquito eggs were hatched in reverse osmosis (RO) water, and 200 larvae were dispersed into rearing trays with 1L of RO water with 2 fish pellets (Hikari). Upon pupation, pupae were transferred to cups and placed inside rearing cages with 20% sucrose *ad libitum*. Adult mosquitoes were maintained on defibrinated calf whole blood (Lampire Biological Laboratories), and eggs were collected in RO water-filled cups placed inside cages two days after blood feeding. Larvae and adult mosquitoes were maintained under standard, controlled insectary conditions in a dedicated double-wide environmental chamber (Percival Scientific) set to 26°C ± 0.5°C, 75% ± 5% relative humidity, and a 14h:10h light:dark diurnal cycle.

### Experimental Design

To determine the effects of constant and fluctuating temperatures on WNV infection and survival in *Cx. tarsalis*, we offered a WNV-infected blood meal to 2,400 4-5 days old adult *Cx. tarsalis* females and an uninfected control bloodmeal to 300 females. Prior to the experimental blood feed, female mosquitoes were deprived of sugar and water for 24 hr. Mosquitoes were blood-fed through a water-jacketed membrane feeder for one hour, after which we randomly distributed 1,800 WNV-exposed and engorged mosquitoes and 180 unexposed blood-fed control mosquitoes into mesh-covered paper cups (30 mosquitoes per cup). Ten infected cups and one control cup were then randomly distributed across six environmental treatments that reflected three mean constant temperatures (22°C, 26°C, and 30°C ± 0.5°C; 75% ± 5% relative humidity; 14h:10h light:dark cycle) and three diurnally fluctuating environments set to a 12°C diurnal temperature range (DTR) around each mean condition (22°C ± 6°C, 26°C ± 6°C, and 30°C± 6°C ± 0.5°C; 75% ± 5% relative humidity; 14h:10h light:dark cycle). Environmental conditions were simulated in upright environmental chambers (Percival Scientific, 36VL) programmed with an asymmetrical minimum-maximum temperature model [52] in which temperature follows a sinusoidal progression during the daytime and a decreasing exponential curve during the night to mimic field-relevant diurnal temperature variations. Mosquitoes were maintained on 20% sucrose *ad libitum* and had their mortality monitored and recorded daily for the duration of the experiment. Two full biological replicates were performed in this study.

### West Nile Virus Culturing and WNV Sampling of Exposed Mosquitoes

For all mosquito infections, we used wild-type WNV (strain FtC-3699, originally isolated from *Culex* spp. mosquito pools collected in Fort Collins, Colorado) and passaged in Vero cells two times. Vero cells (ATCC, CCL-81) were maintained in Dulbecco’s modified Eagle medium (DMEM) supplemented with 5% Fetal Bovine Serum (FBS), and 1% Penicillin/Streptomycin at 37°C and 5% CO_2._ The virus was harvested two days after inoculation and stored at −80°C before titrating. Titers were determined by standard plaque assays on Vero cells as previously described [53] and expressed in plaque-forming units per milliliter (PFU/mL). Mosquitoes were then provided with an infectious bloodmeal spiked with WNV at a final concentration of 1×10^7^ plaque-forming units (PFU)/ml or an uninfected control blood meal. The infectious blood meal was prepared by mixing calf blood with WNV diluted in Dulbecco’s modified Eagle medium (DMEM) at a 1:1 ratio. Every three days, for a total of 30 days post-infection, we force-salivated 24 WNV-exposed mosquitoes per treatment group by immobilizing mosquitoes on ice and removing their legs and wings. Legs and wings were transferred into 250µL DMEM with 20% FBS and 1x antibiotic/antimycotic. We then placed the proboscis of each mosquito into a filtered pipet tip containing 50µL FBS supplemented with 3mM Adenosine triphosphate (ATP) for 30 min on a hot plate at 37°C. After salivation, salivary secretions were then ejected into a screw cap microtube containing 200µL DMEM with 1x antibiotic/antimycotic and mosquito bodies were placed in 250µL of DMEM with 20% FBS and 1x antibiotic/antimycotic. Each tube containing tissues (legs/wings and bodies) was homogenized in a Retsch MM400 grinding Mill at 30Hz for 1:30 min with 2 silica ColiRollers Plating Beads (Sigma-Aldrich) and centrifuged at 14,000 rpm for 1 min at room temperature inside the Biosafety Cabinet. Tubes containing salivary secretions were centrifuged at 14,000 rpm for 1 min inside the Biosafety Cabinet. All samples were stored in a −80°C freezer until use in cytopathic effect (CPE) assays on Vero cells. To measure the proportion of mosquitoes that became infected, had a disseminated infection, and became infectious at each temperature treatment, we tested for the presence/absence of infectious WNV particles in mosquito bodies, legs and wings, and saliva using cytopathic effect (CPE) assays on Vero cells [54].

### Saliva viral load assessment

Viral RNA was extracted from 50µL of mosquito salivary secretions collected in diluent using Quick-RNA Viral Kit (Zymo Research) following the manufacturer’s protocol and eluted in 10µL of nuclease-free water. West Nile virus RNA in saliva was quantified by RT-qPCR using the following forward primer (5’-TCA GCG ATC TCT CCA CCA AAG-3’), reverse primer (5’-GGG TCA GCA CGT TTG TCA TTG-3’), and probe (5’-TGC CCG ACC ATG GGA GAA GCTC-3’) (Grubaugh et al. 2016), and viral loads were determined by comparison to a standard curve generated from serial dilutions of virus of known concentration.

### Statistical Analysis

We ran three sets of statistical analyses investigating the effects of temperature (*T*; 22°C, 26°C, 30°C), daily temperature range (*DTR*; constant or fluctuating), and days post infection (*dpi*; 3, 6, 9, 12, 15, 18, 21, 24, 27, 30) on the probabilities of infection (positive bodies), dissemination (positive legs / wings), and infectiousness (positive saliva); overall viral load in the saliva; and mosquito mortality. First, to estimate the effects of *T*, *DTR*, and *dpi* on the probabilities of infection, dissemination, and infectiousness, we used generalized linear-mixed models (GLMMs) constructed using a binomial distribution and logit link function. *T* and *DTR* were included as categorical factors, while *dpi* was centered by subtracting the mean and scaled by dividing by the standard deviation (SD). To account for differences in WNV infection among mosquito cohorts, we included cohort as a random intercept in each analysis. Because we observe non-monotonic effects of *dpi* on some of our response variables, we incorporated a polynomial function (*dpi^2^*) into our statistical model to accommodate this nonlinearity and tested its interaction with *T* and *DTR*. We compared a series of nine candidate models, ranging from a ‘null model’ with only random effects, a ‘base model’ with only linear effects of each independent variable, to a ‘full model’, including all fixed effects and interactions (Table 4; [55]; package ‘lme4’). Models were ranked using the Akaike information criterion (AIC). To further evaluate model performance, we used the function “*check_model”* from the ‘performance’ package in R, to check for residual dispersion, normality of random effects, homogeneity of variance, and multicollinearity among fixed effects. If models with the lowest AIC showed evidence of multicollinearity or other violations, we selected the most parsimonious model that had both minimized AIC and a satisfactory diagnostic. This procedure ensured that the final model balanced fit with statistical validity. Second, to assess the effects of *T*, *DTR*, and *dpi* on WNV viral loads in mosquito saliva (i.e., WNV genome copies per mL/saliva), we initially fit Gaussian linear mixed-effects models with the same fixed effects structure as our other analyses and mosquito cohort as a random factor. However, the estimated cohort variance was zero, therefore final viral load models were fit as Gaussian linear models without random effects. Candidate viral load models followed the same model selection framework used for infection, dissemination, and infectiousness (Table 4). Finally, to estimate the effects of *T*, *DTR*, WNV exposure, and their interaction on the daily probability of survival, we used a mixed-effects Cox proportional hazards model with *T*, *DTR*, infection status (WNV-exposed or non-exposed), and their interactions as fixed factors, and mosquito cohort as random factor.

**Table 4.** Ten candidate Generalized linear mixed models were selected to explore the effects of temperature, daily temperature range (DTR), days-post infection (DPI) and their interactions on the probability of successful mosquito infection, dissemination, and becoming infectious after being exposed to a WNV-infectious blood meal. Models were run using binomial distribution and logit link. All models included a random intercept for mosquito cohort. For each replicate, the most parsimonious model (M2) is shown in bold.

| Model | Biological hypothesis | Fixed effects |
| --- | --- | --- |
| M0 | The probability of each infection stage is random across batches | None |
| M1 | Temperature, DTR, and time independently influence the probability of each infection stage | $\sim T + dpi + dtr + dpi^2$ |
| **M2** | **Temperature interacts with time to influence the probability of each infection stage, while DTR has a main effect** | **$\sim T * dpi + dtr + dpi^2$** |
| M3 | DTR interacts with time to influence the probability of each infection stage, while temperature has a main effect | $\sim T + dtr * dpi + dpi^2$ |
| M4 | Temperature and DTR interact to influence the probability of each infection stage, while time is independent and non-linear | $\sim T * dtr + dpi + dpi^2$ |
| M5 | Temperature and DTR each independently interact with time to influence the probability of each infection stage, with all pairwise interaction shaping infection dynamics | $\sim T * dtr + T * dpi + dtr * dpi + dpi^2$ |
| M6 | Temperature and DTR jointly modify the effect of time (three-way interaction), while non-linear temporal effects are independent | $\sim T * dtr * dpi + dpi^2$ |
| M7 | Temperature and DTR each modify both linear and quadratic time effects, and also interact to influence overall probabilities | $\sim T * dpi + T * dpi^2 + dtr * dpi + dtr * dpi^2 + T * dtr$ |
| M8 | Temperature and DTR jointly modify both linear and non-linear time effects (Full three-way interactions) | $\sim T * dtr * dpi + T * dtr * dpi^2$ |
| M9 | Temperature modifies both linear and quadratic time effects, while DTR has a main effect | $\sim T * dpi + T * dpi^2 + dtr$ |

Response variable: Mosquito infection status (1 = infected, 0 = uninfected). T = temperature (22°C, 26°C, 30°C), dtr = daily temperature range (constant and fluctuating), dpi = scaled days post-infection, dpi<sup>2</sup> = quadratic term for days post-infection.

### Estimation of vector competence, extrinsic incubation rate, and estimated relative force of infection

Based on previously described methods [48], we estimated how temperature and diurnal temperature range affected WNV vector competence (*bc*) and the extrinsic incubation rate (*EIR*) from the experimental infection data over time, and combined these with variation in the daily probability of survival to quantify the expected availability of infectious mosquitoes through time. Briefly, for each temperature treatment (*T* x *DTR*), we aggregated infection outcomes by day post-infection to estimate the proportion of infectious mosquitoes. We fit a logistic growth model to the proportion of infectious mosquitoes (*Y*) as a function of *dpi* (*t*) for each temperature regime (Eq. 1) using the “nls” package in R [55].

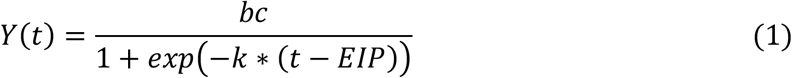

Vector competence is the asymptotic maximum proportion of infectious mosquitoes out of the total exposed for each temperature and diurnal temperature range combination, *k* is the rate parameter, and extrinsic incubation period (*EIP*) is the inflection point (time to reach 50% of maximum *bc*). Because some of our treatments exhibited declines in proportions of infectious at late *dpi* due to mortality, the logistic model was fit to the ascending portion of the time up to the *dpi* at which maximum proportion of infectious was reached. Extrinsic incubation rate (*EIR*) was estimated as the inverse of the *EIP* (*EIR* = 1/*EIP* (*day* − 1)). To estimate the proportion of mosquitoes alive, for each experimental cup, cumulative death counts were converted to daily deaths. For each temperature regime, we fit candidate parametric models (exponential, Weibull, Gompertz, and log-logistic) to the daily survival probability using “flexsurv” package in R [55] and selected the best performing model using AIC.

Finally, we performed an additional calculation to estimate a metric similar to the force of infection following previously described methods [47,48]. Briefly, we multiplied the best fitting non-linear functions describing the daily relationship between the proportion alive and the proportion of infectious mosquitoes for each environmental treatment, resulting in the number of infectious days / environmental condition. We then estimated the area under the curve of the resulting function to calculate the number of mosquitoes that are alive and infectious for a mosquito population of a given size (n=100) over the observed experimental period (30 days).

## Author’s contributions

L.C.M was involved in experimental design, execution, data analysis, and writing this manuscript, D.H. was involved in experimental design and executing this experiment, B.J. helped executing this experiment, A.H. helped execute the RNA extractions and quantitative PCRs, J.S. was involved in maintain the mosquito colony used in the experiments, G.D.E. and C.C.M. led in securing funding for this project, conceptualizing the experimental design, execution, data analysis, and writing of this manuscript. All authors have revised the final version of the manuscript and approved of its contents.

## Acknowledgments

We gratefully thank the members of the Murdock lab for thoughtful comments on the project and manuscript.

## Supporting Information

**Fig. S1. Estimated vector competence of *Culex tarsalis* exposed to West Nile virus.** Relationship between temperature (22°C, 26°C, and 30°C) and estimated vector competence under constant (gray) and fluctuating (blue) temperature regimes. Bars show the estimated maximum proportion of exposed mosquitoes that became infectious and error bars indicate 95% bootstrap confidence intervals.

**Fig. S2. Extrinsic incubation rate of West Nile virus in *Culex tarsalis*.** Relationship between temperature (22°C, 26°C, and 30°C) and extrinsic incubation rate under constant (gray) and fluctuating (blue) temperature regimes. Bars show point estimates for each treatment and error bars indicate 95% bootstrap confidence intervals.

Fig. S3. Effects of temperature, daily temperature range, and days post-infection on WNV saliva viral load (log^10^) in infectious *Culex tarsalis*. A) Model-estimated mean saliva viral loads by temperature, averaged across DTR and dpi. Points represent estimated means, and error bars represent 95% confidence intervals. **B)** Model-estimated viral-load trajectories across dpi under constant and fluctuating temperature regimes. Lines represent predicted means, and shaded ribbons represent 95% confidence intervals. Points show observed median viral loads at each dpi, with error bars representing interquartile ranges.

