## Supplementary figures and images for "The Effects of Mean Temperature and Diurnal Temperature Range on West Nile Virus Transmission"

### Supplemental Figure 1

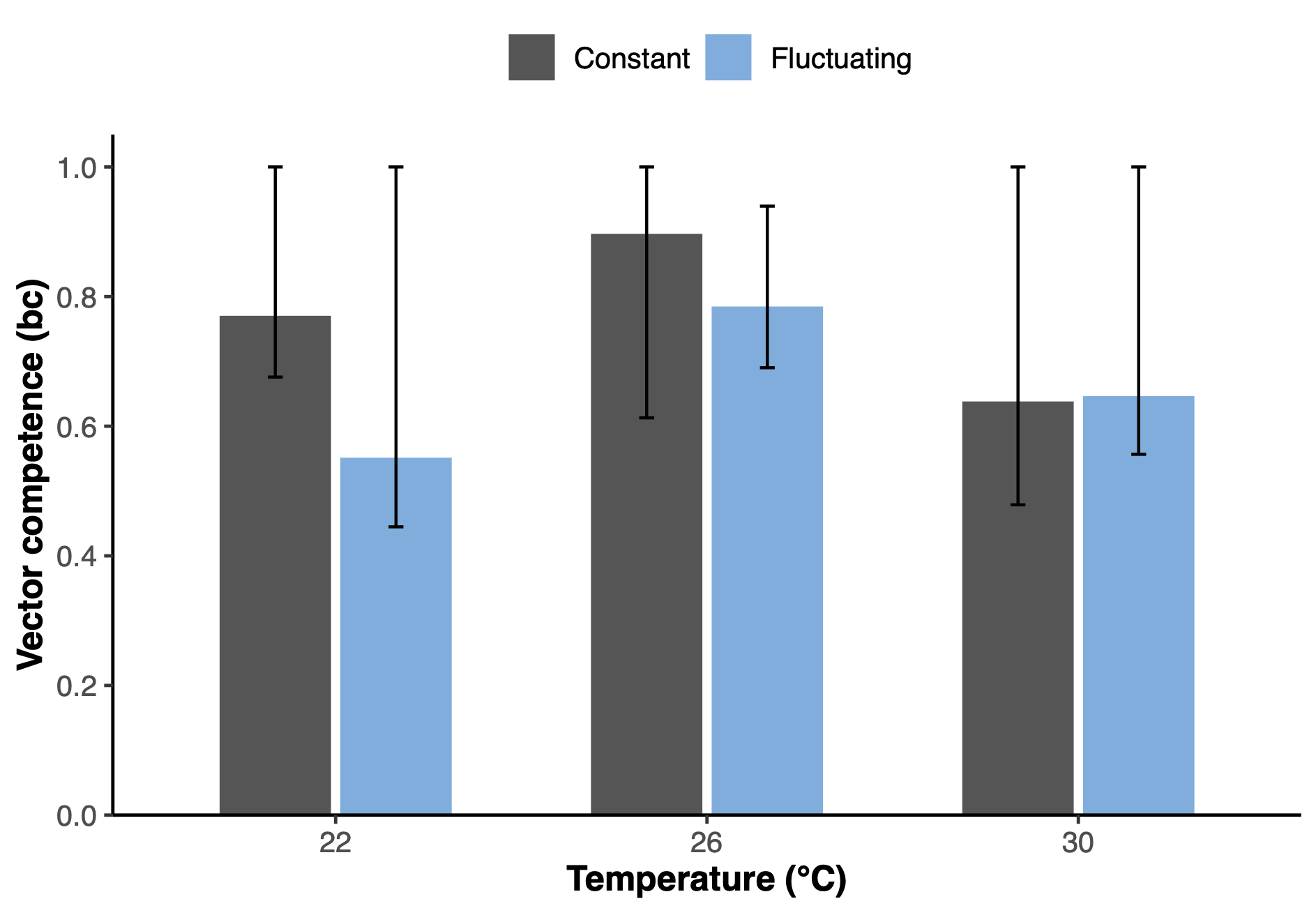

### Supplemental Figure 2

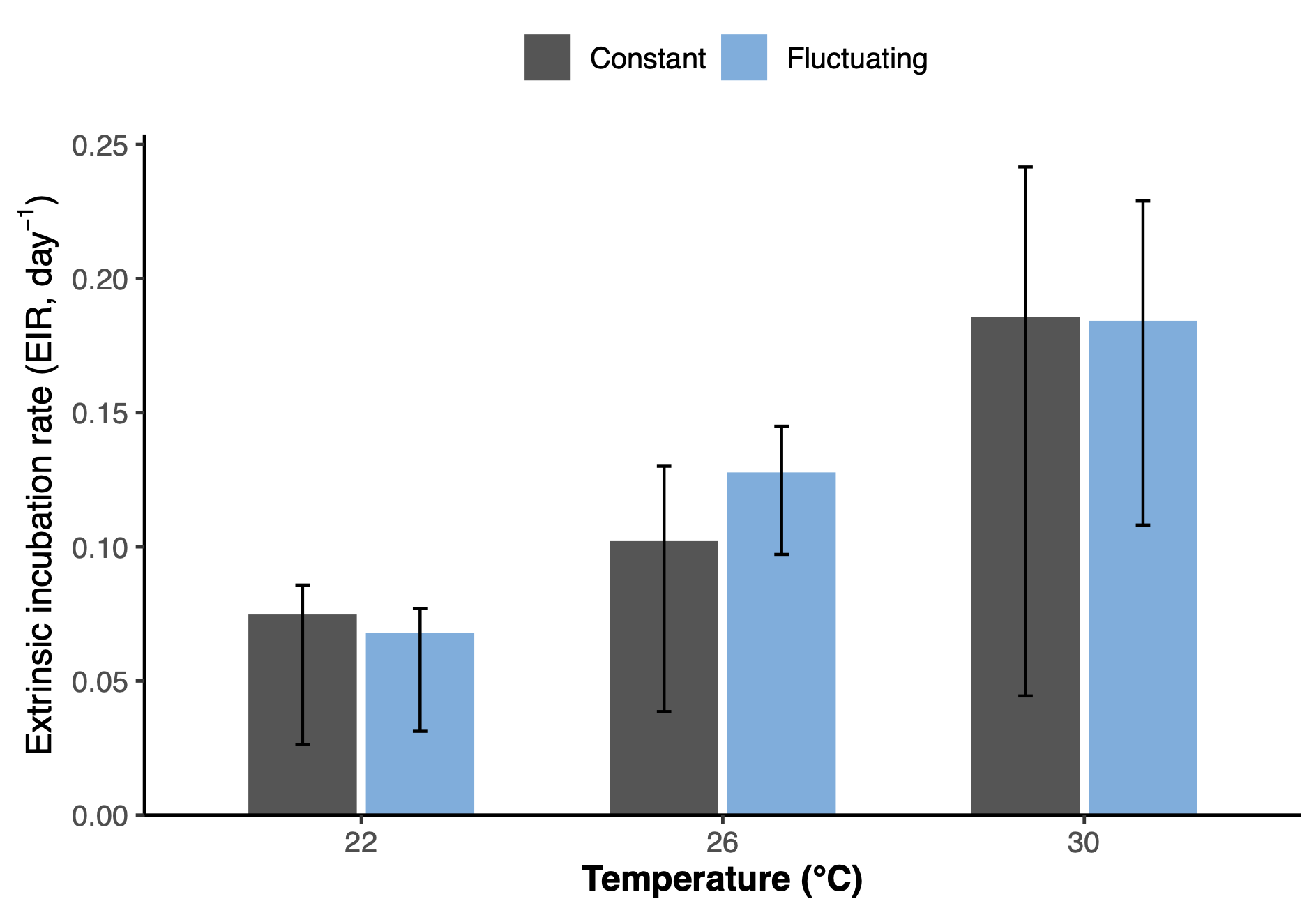

### Supplemental Figure 3

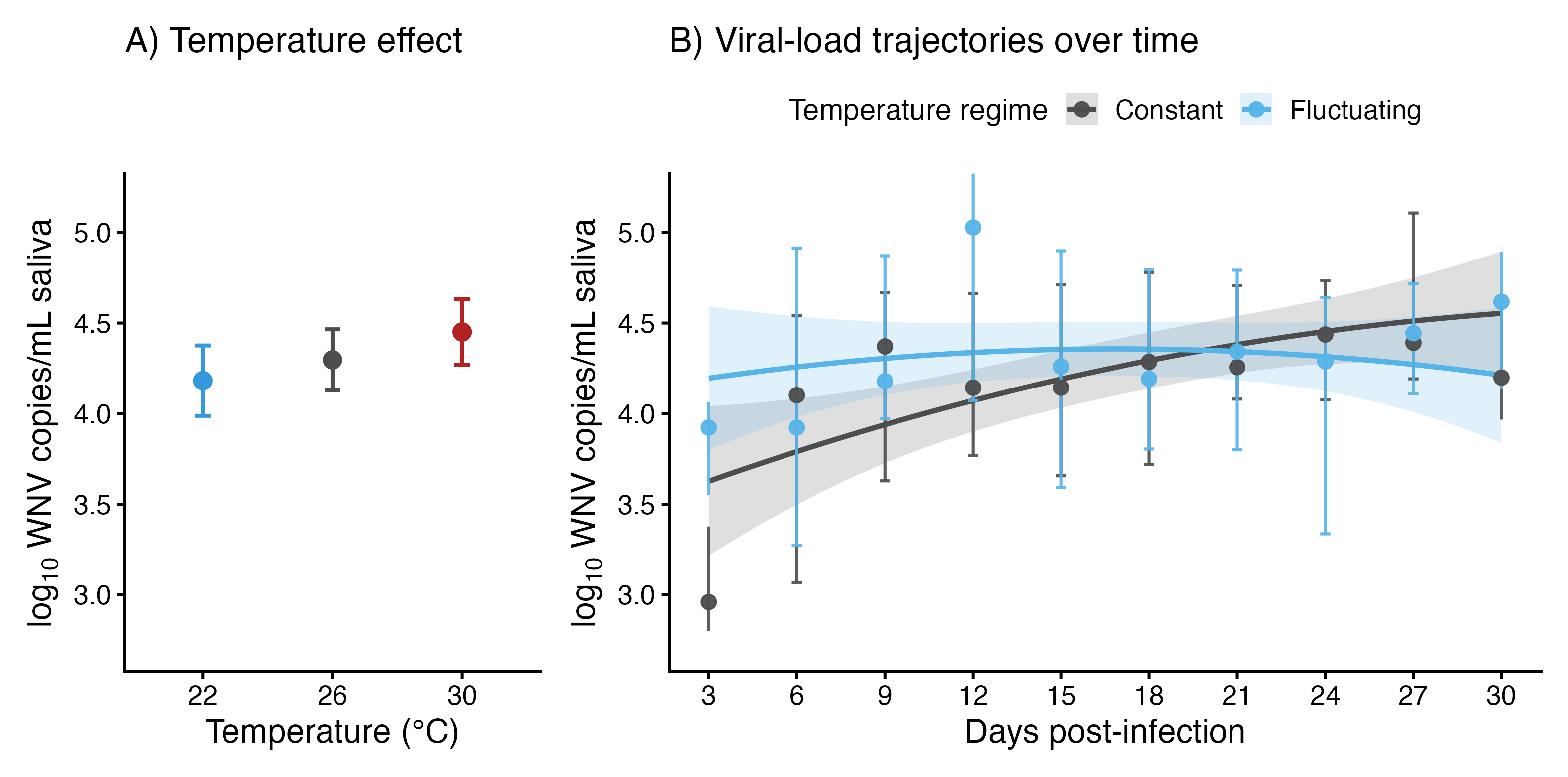
